# Deciphering epigenomic code for cell differentiation using deep learning

**DOI:** 10.1101/449371

**Authors:** Pengyu Ni, Zhengchang Su

## Abstract

Epigenomic markers, such as histone modifications, play important roles in cell fate determination and type maintenance during cell differentiation. Although genomic sequence plays a crucial role in establishing the unique epigenome in each cell type produced during cell differentiation, little is known about the sequence determinants that lead to the unique epigenomes of the cells. Here, using a dataset of six histone markers measured in four human CD4+ T cell types produced at different stages of T cell development, we showed that two types of highly accurate deep convolutional neural networks (CNNs) constructed for each cell type and for each histone marker are a powerful strategy to uncover the sequence determinants of the various histone modification patterns in difference cell types. We found that sequence motifs learned by the CNN models are highly similar to known binding motifs of transcription factors known to play important roles in CD4+ T cell differentiation. Our results suggest that both the unique histone modification patterns in each cell type and the different patterns of the same histone marker in different cell types are determined by a set of motifs with unique combinations. Interestingly, the level of shared few motifs learned in the different cell models reflect the lineage relationships of the cells, while the level of few shared motifs learned in different histone marker models reflect their functional relationships. Furthermore, using these models, we can predict the importance of the learned motifs and their interactions in determining specific histone marker patterns in the cell types.

## INTRODUCTION

Cell differentiation is achieved by remodeling of the same genome that each cell inherits from the zygote, conferring a unique epigenome to each resulting cell type that express a unique set of gene products. Genome remodeling involves methylation of certain cytosine residues in the genomic DNA and various covalent modifications of histones in the nucleosomes (Strahl and Allis 2000). Increasing lines of evidence have suggested that the epigenome in a cell type is established step-wisely though the interplay of genomic sequence, chromatin remodeling systems and environmental cues along the developmental lineage (Juelich et al. 2009; Zhu et al. 2013; Russ et al. 2014; Rodriguez et al. 2017). As the latter two factors are the results of interactions of the products of genomic sequences, the epigenome of a cell type is ultimately determined by the genomic sequence (Thomson et al. 2010; Benveniste et al. 2014; Whitaker et al. 2015). For example, in a recent study, Whitaker and colleagues (Whitaker et al. 2015) have shown that short DNA motifs enriched in the epigenetically modified genomic regions could predict the specific histone modifications in specific cell types using a random forest-based machine-leaning method. However, this method could not discover sequence determinant ab initio because pre-selected motifs were needed to train the models. Therefore, new methods are needed to gain a better understanding of the sequence determinants that specify the unique epigenome of each cell type produced during cell differentiation.

Recent progress in deep-learning has demonstrated that deep convolutional neural networks (CNNs) can achieve very high accuracy in predicting transcription factor (TF) binding affinity (Alipanahi et al. 2015) and epigenetic markers in various cell types (Zhou and Troyanskaya 2015; Kelley et al. 2016; Zeng and Gifford 2017). Unlike traditional neural networks, the kernels in the convolutional layers in a CNN can automatically learn the features of the objects (i.e., the sequence motifs in epigenetically modified regions), and thus the learned featured can provide insight into the underlying mechanisms of the modeled objects. Although efforts have been made to explain the learned motifs in epigenetically modified regions in biological contexts types (Zhou and Troyanskaya 2015; Kelley et al. 2016; Zeng and Gifford 2017), the mixed CNN models employed in these earlier studies lack the power of comparison, limiting their ability to explain the learned motifs for their roles in determining the unique epigenetic modification patterns in different cell types. To overcome these shortcomings, we developed two types highly accurate CNN models to facilitate the explanation of the learned motifs: the cell type model to learn motifs that specify unique patterns of various histone modifications in a cell type, and the histone marker model to learn the motifs that determine different patterns of the same histone marker in different cell types. After demonstrating the superiority of our models over the earlier random-forest-based machine learning method, we evaluated our strategy using a dataset of six histone makers obtained in four human CD4+ T cell types produced at different stages of differentiation (Durek et al. 2016), i.e., the native T (Tn) cells, central memory T (Tcm) cells, T effector memory (Tem) cells and CD4+ terminally differentiated CD45RA+ memory (Temra). The relatively rich knowledge about the regulators and the differentiation process of these T cell subpopulations could facilitate validation of the predictions. Indeed, we found that many sequence motifs learned in the CNN models of both the cell types and histone modifications are highly similar to known binding motifs of transcription factors (TFs) known to play important roles in CD4+ T cell differentiation. Intriguingly, the shared motifs learned in the different cell models support the linear model of CD4+ T cell development, consistent with the earlier results based on the patterns of changes in DNA methylation and DNase accessibility of the genome as well as transcriptomes in the cells (Durek et al. 2016), while the shared motifs learned in different histone modification models reflect the functional relationships of the markers. Furthermore, by computing the impacts of the learned motifs on the prediction of the CNNs, we were able to pinpoint specific roles and interactions of their cognate TFs in determining unique histone modification patterns in different cell types, thereby providing new insights into the underlying mechanisms of histone modifications.

## RESULTS

### The CNN models for cell types are highly accurate and robust for predicting various histone modifications in different cell types

In the genome of a cell type, different loci are modified by the same and/or different chromatin markers. It is the different combinations of these chromatin markers that determine the distinct chromatin states of the genome in different cell types (Ernst and Kellis 2013). To learn the sequence determinants that govern the unique combinations of histone modifications in a cell type, we constructed a CNN model for the cell type for predicting the various histone modifications in its genome. We first evaluated the accuracy of the models using the ENCODE Human embryonic stem cells dataset (Roadmap Epigenomics et al. 2015) that has been studied intensively (Whitaker et al. 2015). To this end, we used 1,038,201, 363,349, 880,462, 1,011,252, and 315,266 histone modification peaks in building the models for the human embryonic stem (H1) cells, trophoblast-like (TBL) cells, mesendoderm (ME) cells, mesenchymal (MSC) cells and neural progenitor (NPC) cells, respectively (Methods). As show in Figure *1*A, the models performed almost equally well for predicting the patterns of six histone makers in the five cell types, with an average AUC ranging from 0.894 (MCS) to 0.933 (H1). However, the models had significantly varying performance for predicting pattern of different histone markers, with the highest performance for H3K4me3 (AUC 0.974), followed by H3K9me3 (AUC 0.938), H3K27me3 (AUC 0.928), H3K36me3 (AUC 0.918), H3K27ac (AUC 0.880) and H3K4me1 (AUC 0.864). We compared our results with those obtained in the earlier study using a random forest-based algorithm on the same dataset (Whitaker et al. 2015). To minimize possible bias of the comparison, we used the AUC values from the original publication. As shown in Figure *1*B, our average AUC across the six markers is 0.917, which is higher than 0.837 achieved by the best full model for the markers in the previous study (Whitaker et al. 2015). Furthermore, our average accuracy for the six markers across the five cell models is 90.6%, which is 11.60 % higher than 79% achieved by the best full model in the earlier study(Whitaker et al. 2015). Compared to the earlier study, we improved the AUCs for all the markers except for H3K4me3 (0.974 vs 0.98). In particular, we increased the AUC for H3K4me1 and H3K9me3 by an average of 0.154 and 0.158, respectively, across the five cell models. The relative performance of the six markers are consistent with the random forest-based method except H3K9me3, which hold the second place in our model while it was ranked fifth in the earlier study.

To evaluate the generality and robustness of our models, we applied them to a dataset collected from four CD4+ T cell types at different stages of the T-cell differentiation (Durek et al. 2016). We used 459,814, 653,272, 978,543 and 2,131,540 histone modification peaks in building the models for the Tn, Tcm, Tem and Temra cells, respectively (Methods). As shown in Figure *3*A, similar to the results from the ENCODE dataset (Figure *1*A), the models also performed almost equally well for predicting the pattern of the six histone markers in the four cell types, with an average accuracy and AUC of 91.53% and 0.916, respectively, which are comparable to those obtained in the ENCODE dataset (accuracy 90.6% and AUC 0.917). However, the models also had varying levels of performance for predicting patterns of different histone markers, with H3K4me3 (average AUC 0.95), H3K27me3 (average AUC 0.932) again being in top three most predictable marker, and H3K27ac (average AUC 0.875) H3K4me1 (average AUC 0.888) again being the two least predictable ones in all the four cell types Figure *4*B. These consistent results in the two datasets from different sources strongly suggest that such difference in the performance of the models might be due to the intrinsic property of the markers, instead of the variation of the quality of datasets. Therefore, our CNN models for cell types are highly accurate and very robust for predicting unique patterns of various histone markers in different cell types.

**Figure 1.**
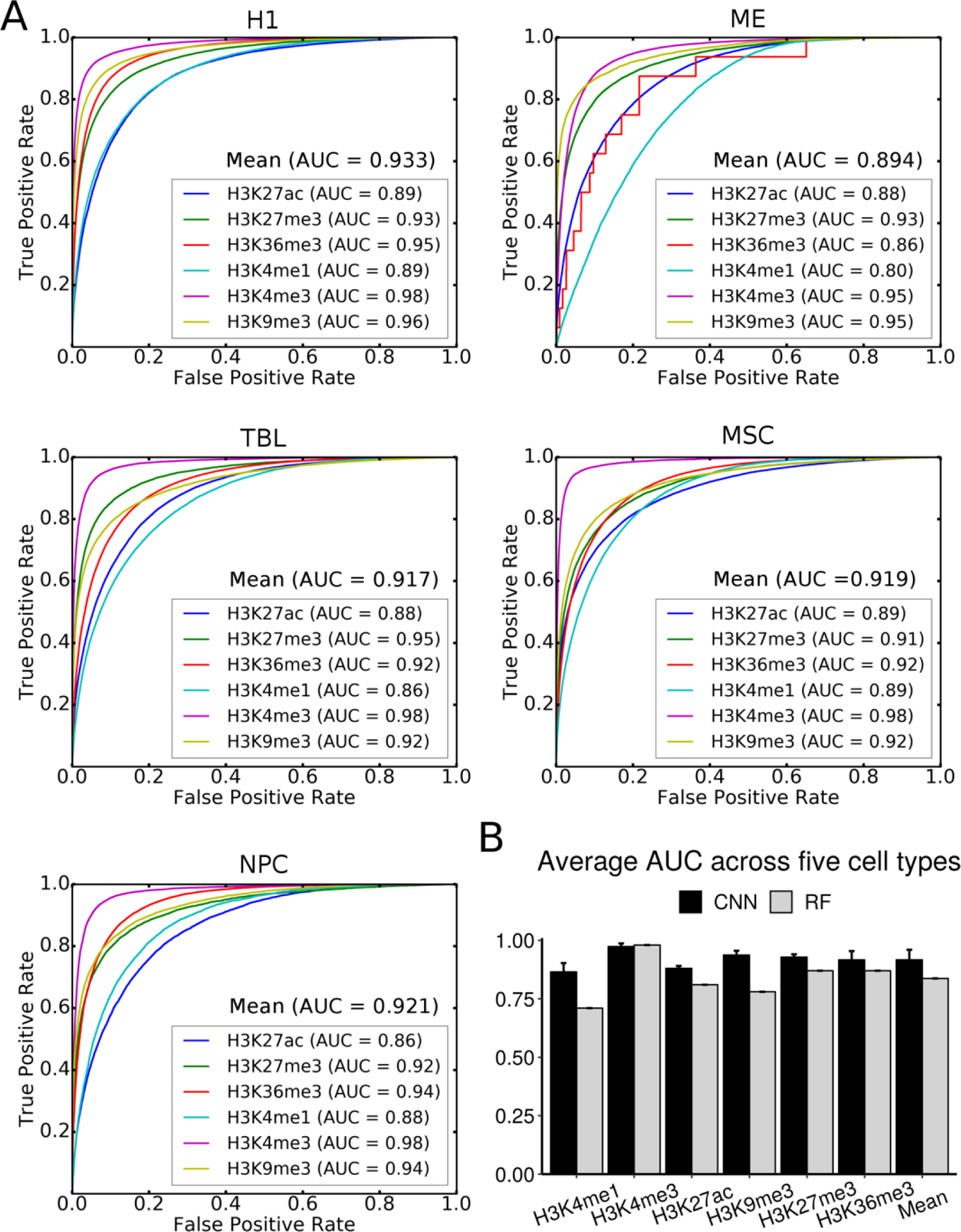
Performance of the CNN models for the five cell types for predicting the six histone markers. A. the ROCs for the histone markers in the H1, ME, TBL, MSC, NPC cells. B. Average AUCs achieved by our CNN models and those obtained by the random forest based models for the marker. The error bars for the random forest based models are not shown due to their unavailability.

**Figure 2.**
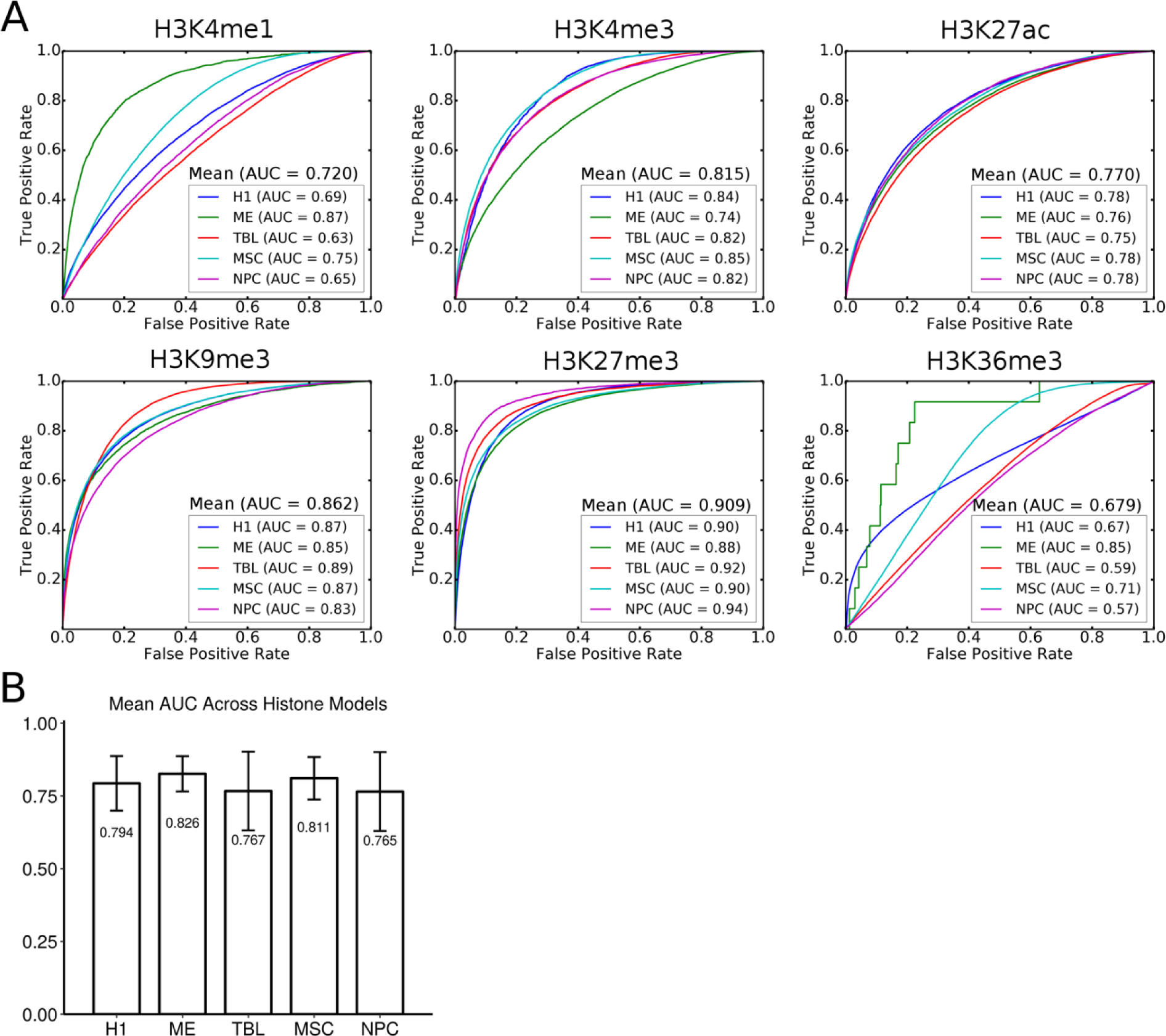
Performance of the CNN models for the six histone makers for predicting the five cell types. A, ROCs of the models for H3K4me1, H3K4me3, H3K9me3, H3K27ac, H3K27me3, H3K36me3, respectively. B. Mean AUC for each cell type across the six histone models.

### The CNN models for histone markers are highly accurate and robust for predicting different cell types based on a histone marker

To reveal the determinants that specify different patterns of the same histone marker in different cell types, we constructed a CNN model for each histone marker for predicting different cell types based on the different pattern of the marker. We also first evaluated the accuracy of the models using the ENCODE Human embryonic stem cells dataset (Roadmap Epigenomics et al. 2015) that has been studied Intensively (Whitaker et al. 2015), and used 332,704, 458,844, 952,615, 185,182, 253,289, 360,040 histone modification peaks in building the models for H3K4me1, H3K9me3, H3K36me3, H3K4me3, H3K27me3 and H3K27ac, respectively (Methods). As show in Figure *2*A, the models had significantly varying performance for predicting the cell types, with H3K27me3 being the highest performer (AUC 0.909), followed by H3K9me3 (AUC 0.862), H3K4me3 (AUC 0.815), H3K4me1 (AUC 0.720), H3K27ac (AUC 0.770) and H3K36me3 (AUC 0.679). Interestingly, gene repression-related H3K27me3 and heterochromatin-related H3K9me3 had better performance than the gene activation-related H3K4me3, H3K4me1, H3K27ac and H3K36me3. However, the average AUCs for predicting the cell types by the five histone markers models were quite similar (Figure *2*B).

To evaluate the generality and robustness of our models, we also applied them to a dataset collected from four CD4+ T cell types (Durek et al. 2016), and employed 227,420, 691,032, 839,057, 867,398, 296,079 and 435,351 histone modification peaks in building the models for H3K27ac, H3K27me3, H3K36me3, H3K4me1, H3K4me3 and H3K9me3, respectively (Methods). As shown in Figure *4*A, similar to the results from the ENCODE dataset (Figure *2*A), the models also had significantly varying performance for predicting the cell types, with AUC ranging from 0.70 for H3K4me1 to 0.95 for H3K27me3. Again, gene repression-related H3K27me3 (AUC 0.95) and heterochromatin-related H3K9me3 (AUC 0.93) had higher performance than the gene activation-related H3K36me3 (AUC 0.87), H3K27ac (AUC 0.85), H3K4me3 (AUC 0.83) and H3K4me1 (AUC 0.70). These consistent results from different datasets from different sources strongly suggest that these two repressive histone markers are more distinctly used in different cell types than the gene activation-related three makers. However, again the average AUCs for predicting cell types by the six histone markers were quite similar (Figure *4*B). Therefore, our CNN models for histone markers are highly accurate and very robust for predicting different cell types based on the pattern of single histone markers.

**Figure 3.**
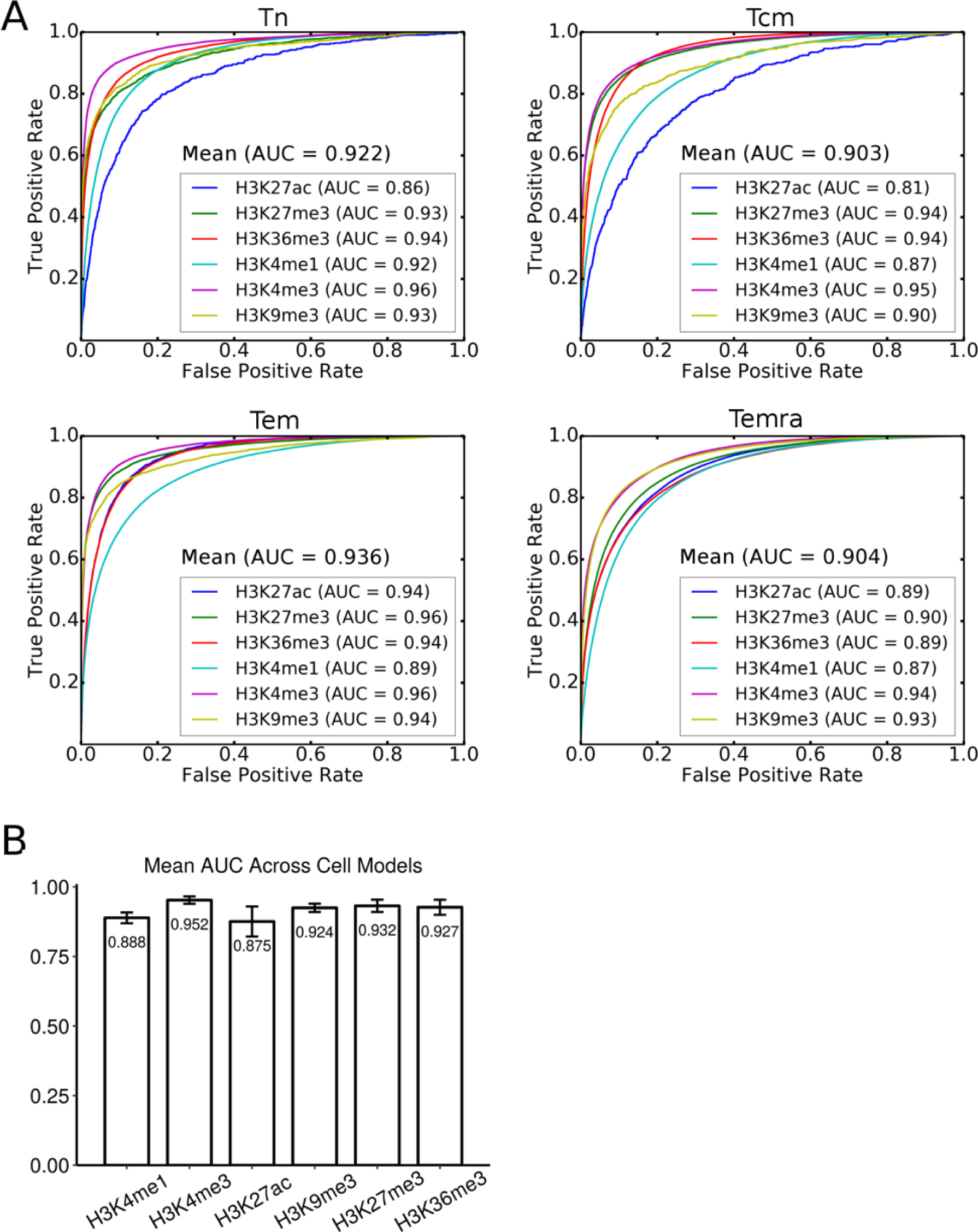
Performance of the CNN models for the four cell types for predicting the six histone markers. A. The ROCs for the histone markers in the native T (Tn) cells, central memory (Tcm) cells, T effector memory(Tem) cells and CD4+ terminally differentiated CD45RA+memory (Temra) cells. B. Mean AUC for each marker across the four cell models.

**Figure 4.**
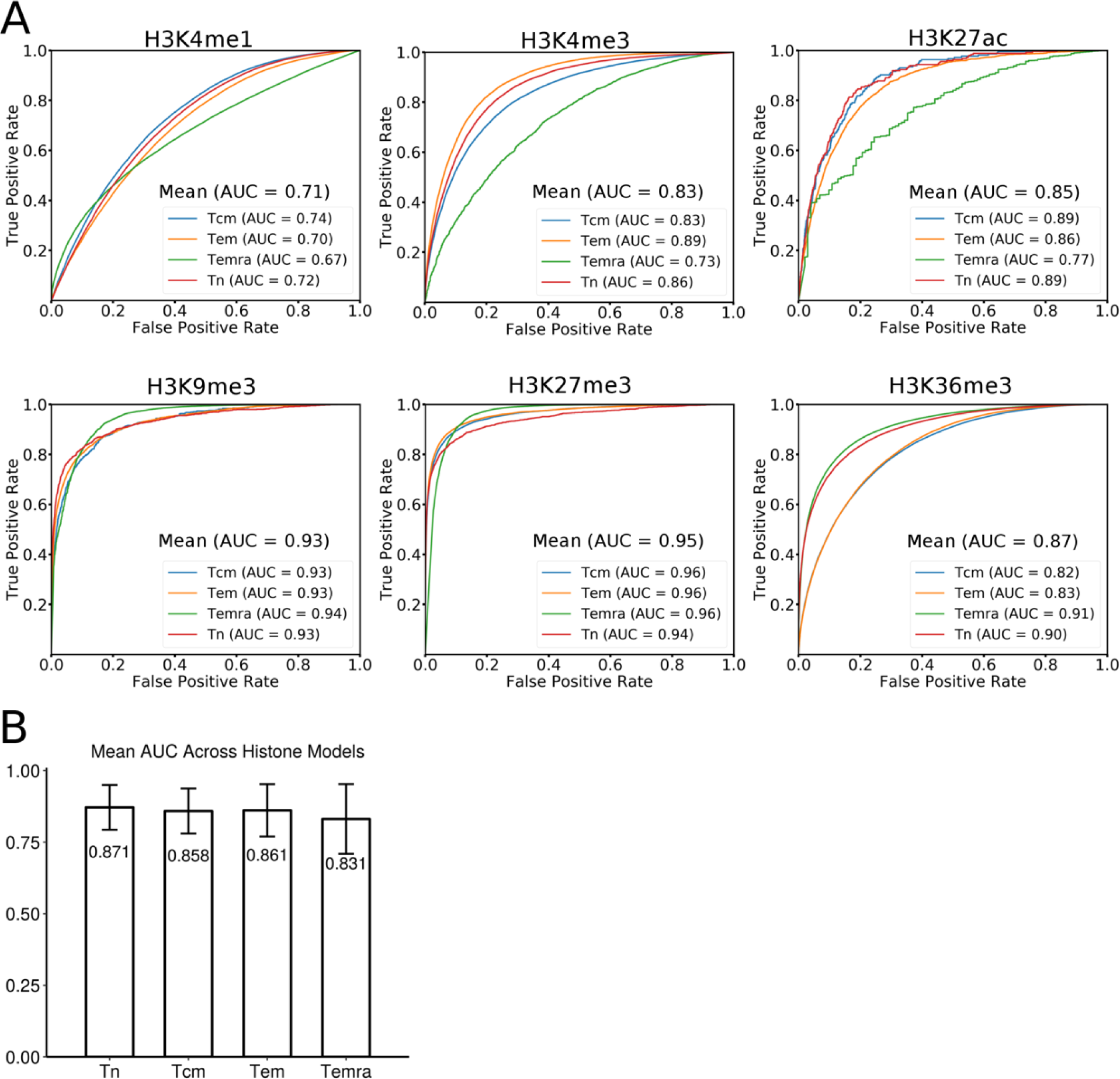
Performance of the CNN models for the six histone makers for predicting the four cell types. A, ROCs of the models for H3K4me1, H3K4me3, H3K9me3, H3K27ac, H3K27me3, H3K36me3, respectively. B. Mean AUC for each cell type across the six histone models.

### Patterns of histone modifications in a cell type as well as different patterns of the same histone marker in different cell types are largely determined by a unique set of motifs

The superior performance of our cell models indicates that the filters in the convolutional layers have largely learned the sequence determinants for specifying the patterns of various histone modifications in each cell type; while the superior performance of our histone marker models suggest that the filters in the convolutional layers have largely learned the sequence determinants for governing different patterns of the same histone marker in different cell types. These results promoted us to reveal these sequence determinants by looking into the filters in the convolutional layers of the CNN models. In particular, we expect that the filters in the first convolutional layer may have learned TF binding motifs enriched in different histone modification peaks. In other words, these filters may correspond to position weight matrices (PWMs) of TF binding motifs. To this end, we constructed motif models for all the filters learned in the first constitutional layers of the models, resulting in 295, 295, 278 and 285 motifs in the Tn, Tcm, Tem and Temra cell models, respectively, and 280, 291, 271, 270, 293, 267 motifs for the H3K27ac, H3K27me3, H3K36me3, H3K4me1, H3K4me3, H3K9me3 marker models, respectively. Some of the motifs learned in different models are highly similar to each other, thus we clustered them according to their similarity (see Methods), resulting in 2,474 clusters. Of these clusters, 203 are formed by more than two learned motifs, and we call each of them a Merged Motif (M-Motif), while the remaining 2,271 are singleton motifs, and we consider each of them as a cell- or marker-specific motif. Interestingly, 113 (4.57%) of these 2,474 unique motifs are shared by at least a cell type model and a histone marker model, suggesting that common sequence determinants might be captured by the two types of models. On the other hand, the remaining 958 and 1,403 motifs are unique to the cell type models and histone marker models, respectively Figure 5A. Thus, besides the common motifs, both the cell type models and marker models captured quite different sets of motifs for predicting the patterns of different histone modifications in cells and the cell types based on single histone markers, respectively. Furthermore, 42 (3.92%) of the 1,071 motifs learned in the cell type models and 68 (4.49%) of the 1,516 motifs learned in the histone marker models are shared by more than two cell models (42/1,071=3.92%) and two maker models (68/1,516=4.49%) Figure 5 B, C, D, E, respectively. However, only two (0.21%) and one (0.10%) motifs are shared by all the four cell type models and all the six histone marker models, respectively. The remaining 1,029 (96.08%) and 1,448 (95.51%) motifs are unique to a single cell type model and a single marker model, respectively. These results suggest that the unique patterns of various histone modifications in each cell type as well as the different patterns of the same histone marker in different cell types are largely determined by quite a unique set of motifs although they may share some common ones. This conclusion agrees with the general understanding about the how the unique epigenomes are established in different cells type by the interplay of TF, chromatin remodeling systems and environment cues (Juelich et al. 2009; Zhu et al. 2013; Russ et al. 2014; Rodriguez et al. 2017).

At an E-value threshold of 0.5, 974 (39.37%) of the 2,474 motifs match known human TF binding motifs in the HOCOMOCO database (Kulakovskiy et al. 2018), and many of them are known to be involved in T cell differentiation (Figure 5F). We described a few examples of them. M-Motif 12 shared by all the cell type models Figure 5F matches that of ETS1 that controls T cell differentiation by regulating the expression of signaling molecules (Li et al. 2000; Wasylyk et al. 2002) in response to external environment stimuli. M-Motif 67 shared by the H3K9me3 and Tem models matches that of ATF2 that is an acetyltransferase (HAT) for histones H2B and H4, playing an essential role in the late-stage activation of T cells (Feuerstein et al. 1996; Kawasaki et al. 2000). Temra-Motif 117 learned in the Temra model matches that of RUNX3, which plays a crucial role in T cell’s differentiation by interacting with master regulators cooperatively(Wong et al. 2011). M-Motif 178 shared by the Tn and H3K4me1 models resembles that of SMAD4 (Figure 5F) that cooperatively regulates interleukin 2 receptor in T cells and balances the differentiation of CD4+T cells (Kim et al. 2005; Malhotra and Kang 2013). H3K27ac-Motif 229 learned in the H3K27ac model matches that of ZN274 that is involved in transcription repression (Valle-García et al. 2016). H3K27me3-Motif 127 learned in the H3K27me3 model resembles that of FOXP1, which is the “naive keeper” for T memory cell differentiation (Hedrick et al. 2012; Durek et al. 2016). These results suggest that at least 39.37% of the learned motifs that match known ones are likely to be authentic motifs of the cognate TFs.

### Motifs learned in the cell type models reflect the lineage of the cells

It is now well established that along the lineage of cell differentiation, the epigenomes of cells undergo step-wise changes through the regulation of a specific set of both common and unique TFs in the derived intermediate and terminal cell types (Juelich et al. 2009; Zhu et al. 2013; Russ et al. 2014; Crompton et al. 2016; He et al. 2016; Rodriguez et al. 2017). Cells in adjacent differentiation stages possess more similar epigenomes (Alipanahi et al. 2015), presumably because they share more TFs than those that are distal from each other along the lineage of differentiation. To see whether this is reflected in the motifs learned by the cell type models, we hierarchically clustered the cell types based on the similarity of learned motif profiles in the cell type models. As shown in Figure 5G, Tn branches earliest in the tree while the three memory/effector T cell types forms a clade, indicating that Tn is most distinct from the more developed cell types as generally believed. Tem and Temra forms a clade, indicating that they are more similar to each other than to Tcm, which is in agreement with early observations(Henson et al. 2012). These results suggest a linear lineage model of the development of these cells: Tn → Tcm → Tem → Temra, which is in line with the results derived based on changes in the DNA methylation, gene expression and DNAase accessibility in the cells (Durek et al. 2016). Therefore, the sequence motifs learned in the cell type models indeed reflect the lineage relationships of the cells. It is highly likely that the unique motifs to a cell model account for the distinction of the cell type from the other cell types, while the shared motifs are responsible the shared features of linearly closely-relate cell types.

### Motifs learned in histone marker models reflects functional relationships of the markers

It is well-known that certain types of sequences can be co-modified by different histone markers, while other types of sequences tend to be exclusively modified by a specific marker (Wang and Willard 2012). To see whether such co-modifications and exclusiveness of the markers are reflected by the learned motifs in the histone marker models, we hierarchically clustered of the histone markers based on the similarity of the learned motif profiles. As shown in Figure 5G, H3K4me1 and H3K27ac form a group, which is consistent with the fact that they co-mark active enhancers, thus the respective modification systems might be recruited by some common motifs or similar mechanism. On the other hand, H3K9me3, H3K27me3, K3K36m3 and H3K4me3 form a singleton group by themselves, which is consistent with the facts that they exclusively mark DNA domains with different epigenomic states in humans (Ho et al. 2014). For instance, H3K9me3 marks heterochromatins, H3K27me3 labels polycomb-associated domains, K3K36m3 marks transcribed gene body, and H3K4me3 labels is active promoters. Therefore, the learned motifs in the histone marker models indeed reflects the known functional relationships of the markers.

**Figure 5.**
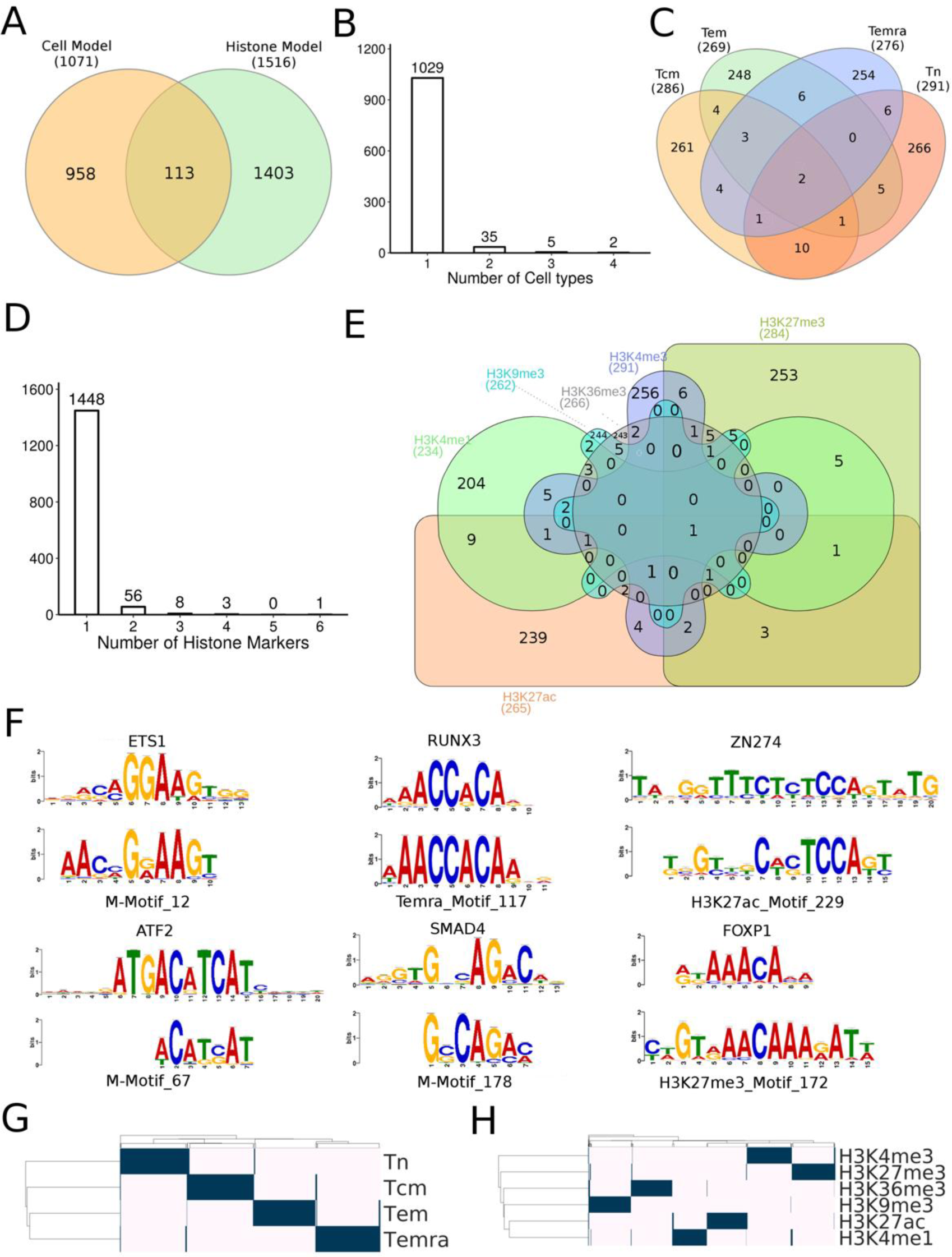
Known and novel TF binding motifs learned in the first convolution layer of the CNN models. A. Overlap of motifs learned in the cell models and histone marker models. B. Number of learned motifs shared by different number of cell models. C. Venn diagram showing the number of learned motifs shared by the cell models. D. Number of learned motifs shared by different number of histone marker models. E. Venn diagram showing the number of learned motifs sheared by the histone marker models. F. Examples of learned motifs matching known motifs involved in T cell functions. G. Hierarchical two-way clustering of the cells based on the similarity of the learned motifs profiles in the models H. Hierarchical two-way clustering of the histone makers based on the similarity of the learned motifs profiles in the models. Venn diagrams were drew using INteractiVenn (Heberle et al. 2015)

### The learned motifs have varying impacts on the predictions of the models

To evaluate the contribution of a learned motif to the prediction of a model, we nullified the motif and then calculated its impact score (see Methods). The impact scores of the motifs learned in both the cell type models (Figure 6A) and the histone marker models (Figure 6B) have a bell-shape distributions with different extent of right skewness. These results suggest that most learned motifs have intermediate impacts, while a small portion have large impacts on predicting patterns of different histone markers in a cell type or different cell types based on single histone markers. The impacts of the motifs learned in both the cell type models (Figure 6A) and histone marker models (Figure 6B) do not significantly correlate with their information contents, suggesting that only few positions of the motifs have a strong predictive power, which is consistent with the general understanding about the mechanisms of TF-DNA interactions. The learned motifs that do not match known motifs have similar impact scores to those matching known motifs (Figure 6A and B), indicating that they are equally likely to be true motifs TFs, and the unmatched ones are likely to be novel motifs of unknown TFs.

Interestingly, the impacts of motifs learned in Tn, Tcm, Tem cell models increased along the proposed linear cell lineage, and then decreased in the Temra cell model (Figure 6C). These results suggest that the functions of learned motifs become more and more specific in determining the patterns of various histone modifications in the cells along the differentiation lineage Tn → Tcm → Tem, and then somehow become less specific in Temra. Furthermore, the impact scores of motifs learned in the six histone marker models are also significantly different from one another (Figure 6D). Specifically, motifs learned in the models of H3K4me1, H3K27ac and H3K4me3 that mark active enhancers and promoters have the lowest impact scores, while those learned in the models of H3K9me3 and H3K27me3 that are associated with repression regions have the moderate impact scores, and those learned in the model of H3K36me3 that marks actively transcribed regions have the highest impact scores (Figure 6B). These results suggest that the motifs specifying histone modifications in actively transcribed regions have the highest specificity, followed by those for determining histone modifications in repression regions, active promoters and enhancers regions.

**Figure 6.**
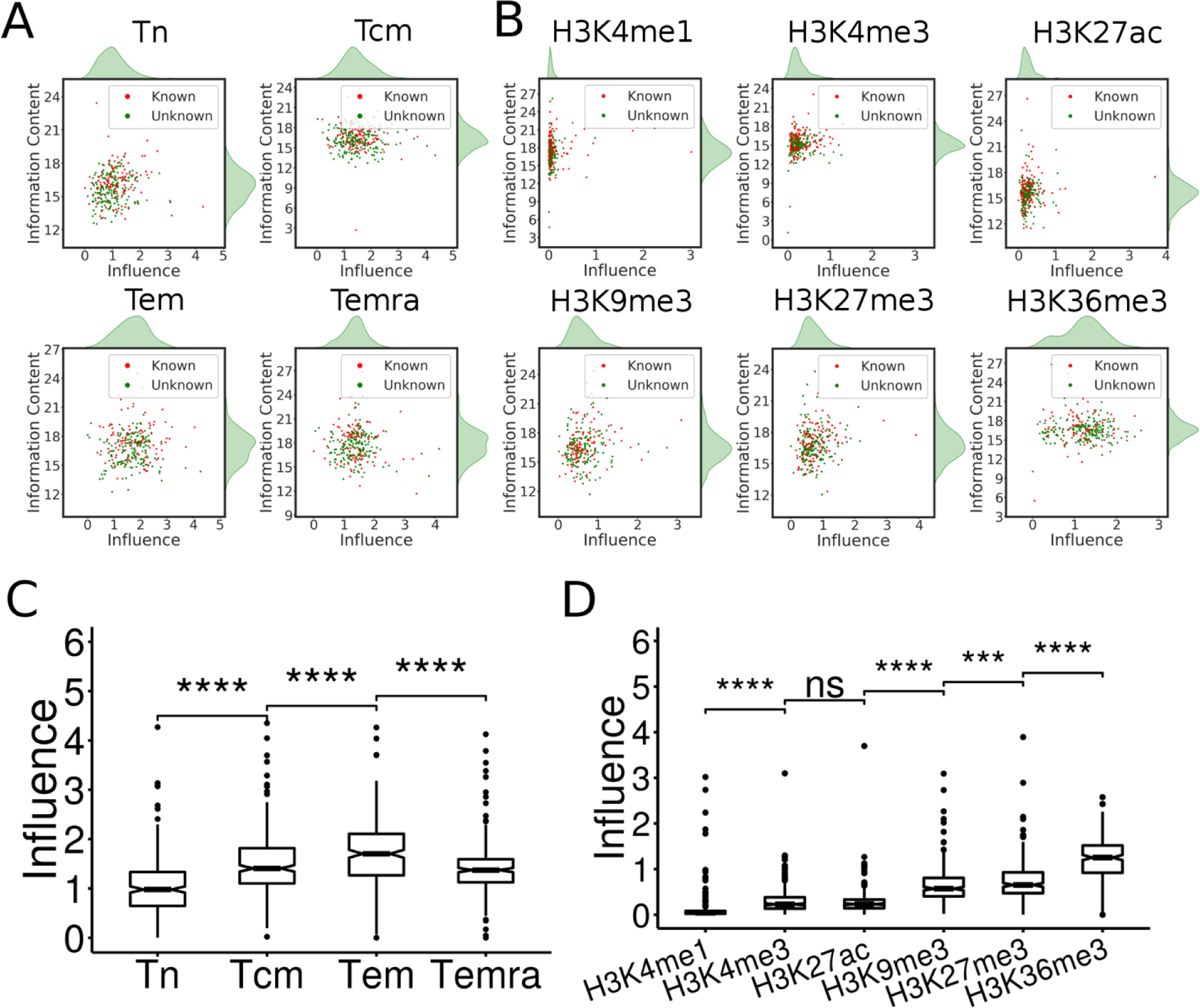
Distribution of the impact scores of learned motifs in cell and histone marker models. A, B. Relationship between the impact scores and information contents of motifs learned in the four cell models and six histone models, respectively. C, D. One-Way ANOVA analysis of the impact scores of the motifs learned in cells models and histone models using the Wilcoxon test, respectively

### The motifs learned in a cell type model have highly variable impacts on different histone markers

An important question in epigenomics study is to understand how different histone markers are placed at specific domains of the genome in a cell type. Our cell models might provide an easy way to address this question by simply finding out the learned motifs that impose a high impact on the prediction of each histone marker modification in the models. More specifically, we computed an impact score of each learned motif on each histone marker in a cell type model. Shown in (Figure 7) are the results for the learned motifs that are ranked top 100 for their impacts on predicting at least one histone marker modification in the four cell type models. Clearly, the motifs learned in each cell type model have highly variable impacts on different histone markers. For example, in all the four cell types, H3K36me3 and H3K27me3 are highly impacted by a large number of the learned motifs, while H3K4me3 is only highly impacted by a few learned motifs, such as Tn-26:FOXD1, Tn-106:HXB4, TN-21 and Tn- 294 in Tn (Figure 7). H3K27ac is highly impacted by a large number of learned motifs in Tcm, but is highly impacted by only a few learned motifs in Tn, Tem and Temra. H3K4me1 is highly impacted by a larger number of learned motifs in Tcm, Tem and Temra, but is highly impacted by a few learned motifs in Tn. H3K9me3 is highly impacted by an intermediate number of learned motifs in all the four cell types. Moreover, in all the four cell models, only a few learned motifs have high impacts on all the histone markers, while most motifs have a high impact only on 1-3 histone markers (Figure 7). For instance, in Tn model, only motifs Tn-26:FOXD1, Tn-106:HXB4, Tn-21 and Tn-294 have high impacts on all the six histone markers, while most of other motifs have high impacts only on one or two histone markers. Thus each histone marker is impacted by a unique combination of motifs that may have impacts on more than two histone markers. These results suggest that the cognate TFs of most learned motifs exerting more specific impacts on one or two histone markers might play crucial roles in specifying the unique patterns of different histone markers in the cell type, while the cognate TFs of a few learned motifs having high impacts on multiple histone markers might be involved in the establishment of multiple histone markers, probably by playing roles in the common mechanisms of different histone modifications such as opening up of DNA domains. 19

**Figure 7.**
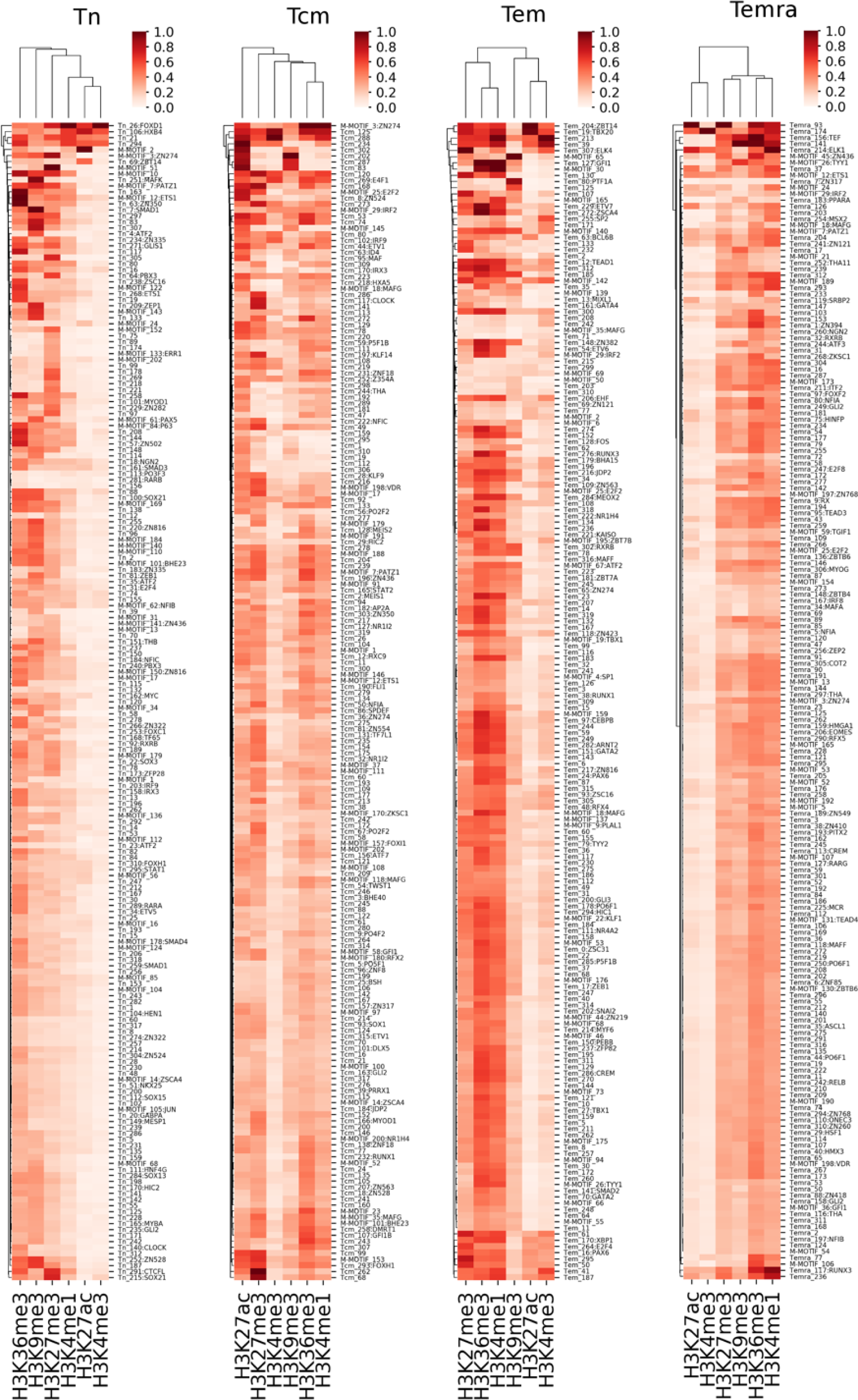
Impacts of the learned motifs on the prediction of each histone marker in the cell models.

### The motifs learned in a histone marker model have highly variable impacts on different cell types

Another important question in epigenomics study is to understand how the same histone marker is differentially placed in the genomes of different cell types. Our histone marker models might provide a convenient way to tackle this question by simply identifying the learned motifs that impose a high impact on the prediction of each cell type in the models. More specifically, we calculated an impact score of each learned motif on the prediction of each cell type in a histone marker model. Shown in Figure 8 are the result of motifs that are ranked top 100 for their impacts on predicting at least one cell type in the six histone models. Interestingly, motifs learned in each the histone marker model have highly variable impacts on different cell types. In the H3K4me1 model, most of the learned motif have similarly small impacts on all the four cell types, only few have high impacts on at least one cell type. However, the latter set of motifs exert high impacts only on one or two cell types with the exception that motif H3K4me1-236:HXC10 have high impacts on all the four cell types. Thus it seems that H3K4me1 in each cell type is specified by a small set of motifs with unique combinations. In both the H3K4me3 and H3K27ac models, most of the learned motif have similarly small impacts on the Tem, Tcm and Tn cell types, only few have high impacts on at least one of these three the cells types, suggesting that these two histone modifications are specified by a small set of motifs with unique combination in these three cell types. However, most of the motifs learned in the H3K4me1 and H3K27ac models impose high impacts on the Temra cells, suggesting that these cells might have more complex H3K4me3 and H3K27ac modifications than do the other three cell types, which is in line with fact that Temra is the terminally differentiated cells with more activated enhancers and promoters. In the H3K9me3 and H3K27me3 models, each cell type is impacted by a large number of learned motifs with few having high impacts on more than three cell types, suggesting these two histone modifications in each cell type are specified by a large set of motifs with unique combinations. This result might be related to the functions of H3K9me3 that marks heterochromatins and of H3K27me3 that labels polycomb-associated domains. In the H3K36me3 model, the numbers of learned motifs having high impacts in the cells increase along their linear lineage: Tn → Tcm → Tem → Temra. Each cell type is highly impacted by a large number of the learned motifs impacting adjacent cells along the lineage. These results reflect the similarity of the transcriptomes of these cell types (Whitaker et al. 2015), and thus is in excellent agreement with the function of H3K36me3 that marks actively transcribed genes. Taken together, the cognate TFs of few learned motifs having high impacts on multiple cell types might account for the similar patterns of a histone marker and the common mechanisms of the histone modification in different cell types, while the cognate TFs of the sets of motifs having more specific impacts might play crucial roles in specifying the different patterns of the histone modification in different cell types.

**Figure 8.**
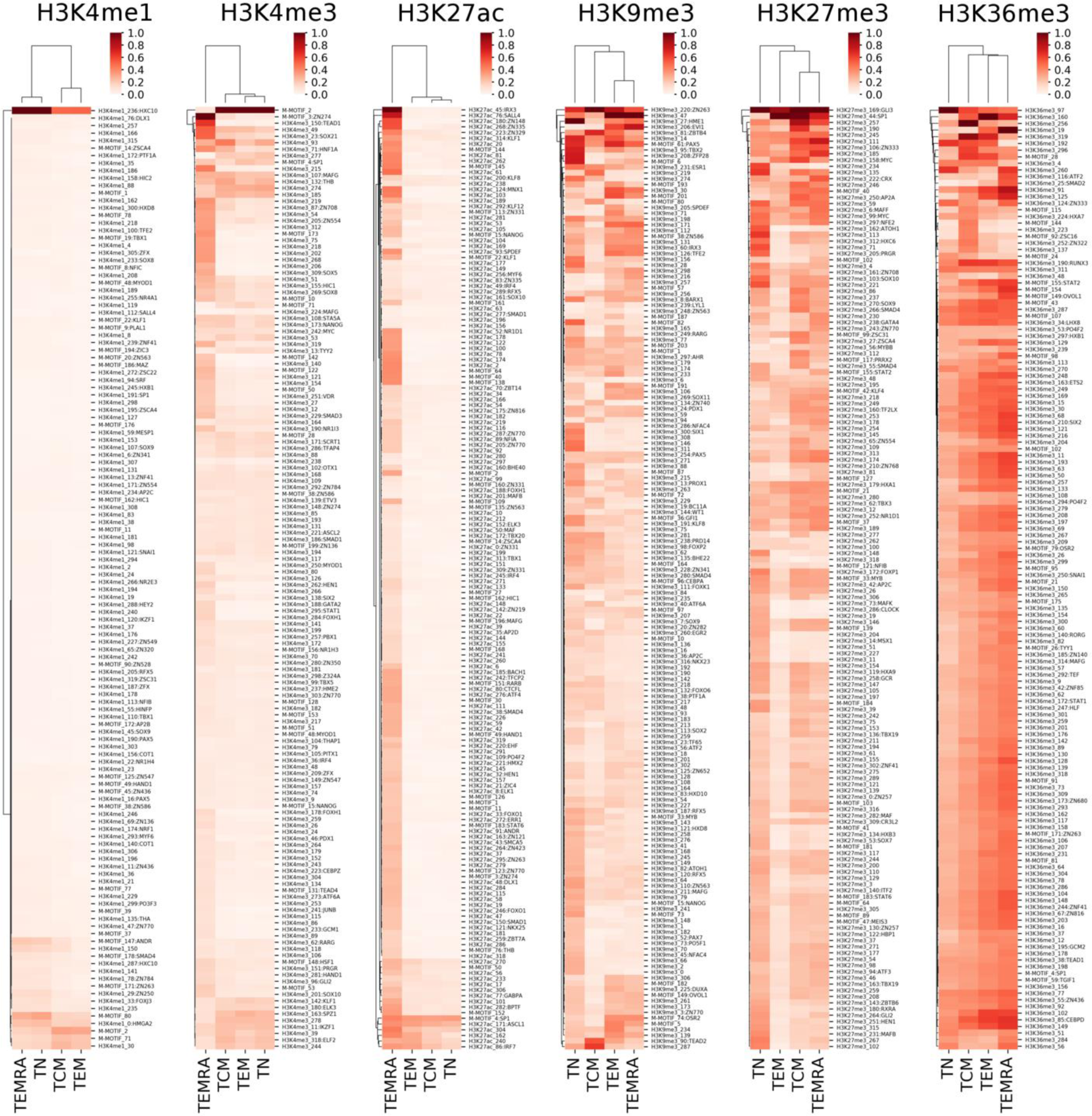
Impacts of the learned motifs on the prediction of each cell type in the histone models.

### Conserved learned motifs tend to have higher impacts on the predictions

We also examined the relationships between the impact scores and the conservation levels of the motifs learned in the cell and histone marker models. As shown in Figure 9A, there is moderate positive correlation between the impact scores and the conservation levels of motifs learned in all the cell models (Tn: r=0.15, p=0.011; Tcm: r=0.11, p=0.052; Tem: r=0.079, p=0.19; and Temra: r=0.17, p=0.003), though with varying levels of significance. Moreover, as shown in Figure 9B, there is significantly moderate positive correlation between the impact scores and the conservation levels of motifs learned in the models of the four activation-related histone markers H3K4me1 (r=0.43, p=2.3e-13), H3K4me3 (r=0.17, p=0.0043), H3K27ac (r=0.35, p=2.6e-9) and H3K36me3 (r=0.23, p=0.00016). However, there is negative or no significant correlation between the impact scores and the conservation levels of motifs learned in the models of the two repression-related markers H3K9me3 (r=-0.13, p=0.036) and H3K27me3 (r=0.063, p=0.29). These results indicate that more conserved motifs learned in either the cell or histone marker models generally have higher impacts on the respective predictions than less conserved ones, with the exception that rapidly evolving motifs in the H3K9me3 marked peaks (heterochromatins) tend to have higher impacts on the prediction of cell types than more conserved ones. These observations are in line with the general understanding of the evolution of DNA sequences that functionally important sequences tend to be either more conserved due to purifying selection or evolved more rapidly due to positive selection. Thus these results also further corroborate our predicted motifs.

Interestingly, motifs learned in the Tn and Tcm models tend to be more conserved than those learned in the Tem and Temra models, and and the motifs learned in the Temra model are least conserved. Thus, there is a trend that the more differentiated the cells, the less conserved the motifs learned from the corresponding models, suggesting that more conserved mechanisms might be used to in the cells at the earlier stages of differentiation to specify their histone modification patterns than in cells in the later stages of differentiation. This conclusion is consistent with the general understanding about the development of animals from cellular to organ levels (Gilbert 2000). Moreover, motifs learned in the models of gene activation-related markers H3K4me3, H3K27ac, H3K4me1 and H3K36me3 are more conserved than those learned in the models of repression-related markers H3K9me3 and H3K27me3. This result suggests that more conserved mechanisms might be used to specify the patterns of the four activation-related markers than those used to govern the patterns of the two repression-related markers.

**Figure 9.**
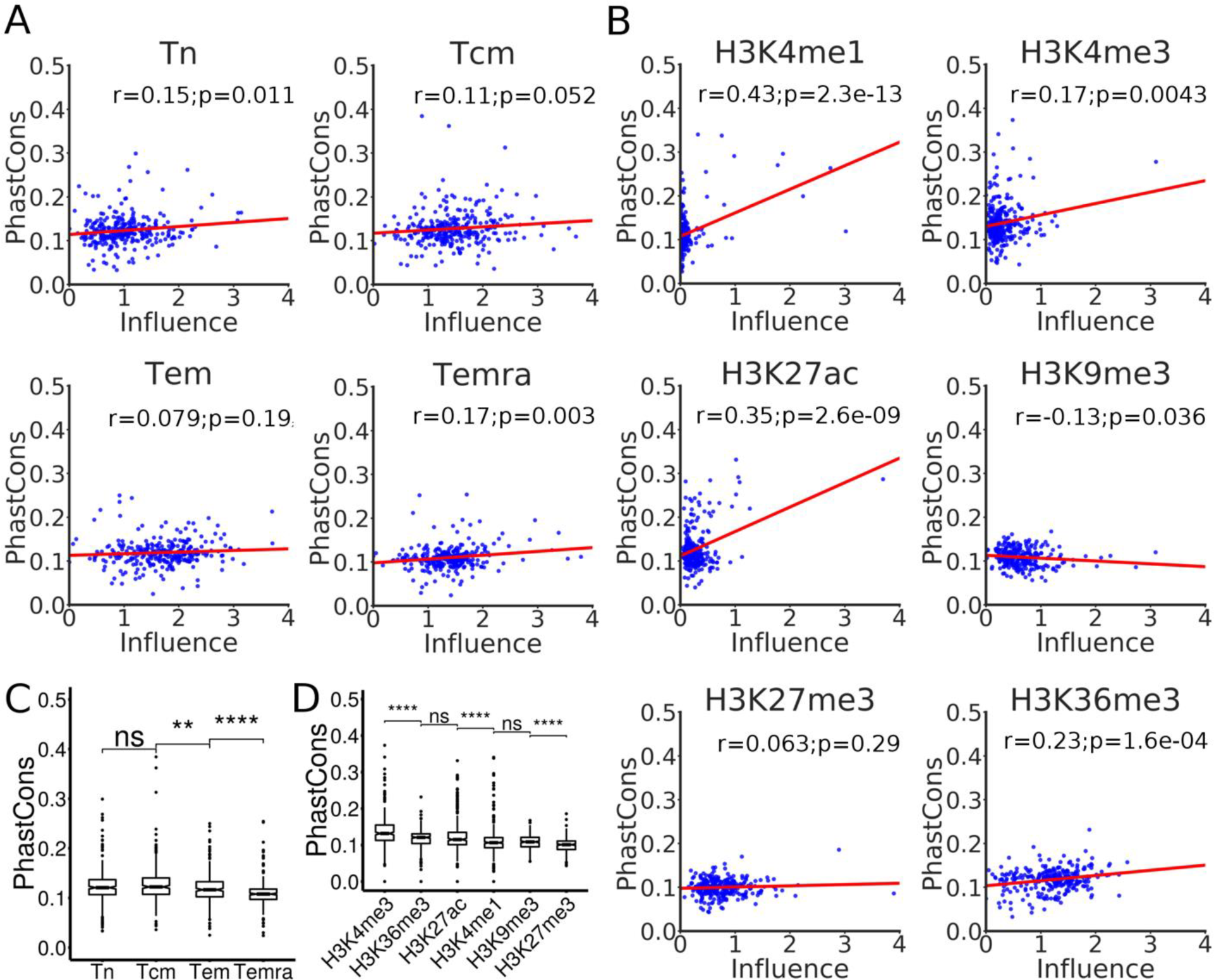
*Relationship between the impact scores and PhastCons scores of the learned motifs. A, B. Relationship between the* impact scores and PhastCons scores of the learned motifs in the cell models and in the histone marker models, respectively. The red line is the linear regression between the impact scores and PhastCons scores. C, D. One-way ANOVA analysis of the PhastCons scores of the motifs learned in cells models and histone models using the Wilcoxon test, respectively.

### The CNN models can predict cooperative TFs for specifying histone modifications in cells

To see if the models can be used to identify cooperative TFs that define the histone modification patterns in the cells, we quantified the interactions between each pair of learned motifs using a linear regression model where a positive or negative interaction coefficient indicates positive or negative interaction (Methods). To reduce the computational time, we only focused on the top 50 of learned unique motifs with the highest impact scores for the cell models and histone marker models. Shown in Figure 10 are the results for the model of Temra. Clearly, there are different patterns of positive and negative interactions between the learned motifs for predicting different histone makers. Interestingly, the motifs can be clustered into groups based on the patterns of their interactions in predicting the histone modifications. For example, in the case of predicting H3K4me1 modifications, the motifs matching those of RUNX3, ETS1 and PATZ1 form a group with positive interactions among them; motifs matching those of EOMES, NFIA, ELK1, HINFP and ITF2 form a group with many other novel motifs with largely positive interactions among them; motifs matching those of TEAD3, ZN121, HMGA1, ZN436, GLI1, ZN274, COT2, RX, TEF, ZN394 and TYY1 form a group with many other novel motifs with largely negative interactions among them. Many of the predicted interactions are supported by experimental evidences. For example, we predicted ITF2 (also named T cell specific transcription factor 4 (TCF4)) had significant interactions with ETS1 for predicting histone modifications H3K27ac (γ=1.27, p=3.69e-65), H3K27me3 (γ=0.18, p=0.01), H3K36me3 (γ=0.21, p=0.00077), H3K4me3 (γ=1.15, p=8.54e-57) and H3K9me3 (γ=-0.39, p=6.70e-06), and in agreement with these predictions, it has been shown that ITF2 might involve in histone acetyltransferase CBP recruitment by interacting with ETS1 to form a complex (Tushir and D'Souza-Schorey 2007). Furthermore, we predicted that ITF2 had a positive interaction with RUNX3 for determining histone modifications H3K27ac (γ=1.40, p=4.29e-49), H3K27me3 (γ=-0.20, p=0.013), H3K36me3 (γ=-0.59, p=6.40e-25), H3K4me1 (γ=0.32, p=8.91e-05), H3K4me3 (γ=-1.13, p=4.00e-40), and H3K9me3 (γ=-0.18, p=0.03), which is in line with the earlier finding that RUNX3 interact with ITF2 (TCF4) forms a ternary complex, which involved in attenuating Wnt signaling activity (Ito et al. 2008). The predicted interactions between known and unknown motifs as well as between unknown motifs are likely to be novel interactions, in particular those with strong and highly significant interactions, such as the interactions for predicting H3K27ac, between GLI2 and Temra 146 (γ=2.137, p=4.95e-43), between TEAD3 and Temra 54 (γ=1.97, p=4.43e-50), and between Temra 141 and Temra 146 (γ=1.99, p=4.41e-43), etc. Similar patterns of interactions were obtained in the models of the other three cell types (Supplementary Figures 1-3).

Shown in Figure 11 are the results for the H3K4me1 model. Again, there are distinct patterns of positive and negative interactions between the motifs for predicting different cell types. As in the cases of cell models, the motifs can be clustered into groups based on the patterns of their interactions for predicting the cell types. For instance, in the case of predicting the Tn cells, the putative novel motifs M- Motif-71 and H3K4me1-30 form a group with a negative interaction; motifs matching those of HIC2, HXD2, TFE2, ZN547, HAND1, COT1, SMAD4, TBX1, ANDR, ZN263, THA, ZN784, ZSCA4, ZN436, PTF1A and ZN770 form a group with many other novel motifs with largely positive interactions among them; motifs matching those of HXC10, PO3F3, POXJ3, HMGA2, HXC10, DLX1 and ZN250 form a group with many other novel motifs with largely negative interactions among them. Many of the predicted interactions are supported by experimental evidences. For example, we predicted that TFE2 interact with HAND1 for predicting Tn (γ=5.38, p=1.84e-137), Tcm (γ=4.00, p=6.94e-115), Tem (γ=2.82, p=1e-70) in Temra (γ=-7.61, p=1.97e-45), while it is has been reported that TFE2 (also named E47) directly interacts with HAND1 (Morin et al. 2005). We predicted that SMAD4 interacts with ANDR for predicting Tn (γ=2.91, p=2.68e-47), Tcm (γ=3.49, p=1.86e-77), Tem (γ=2.99, p=3.47e-79) and Temra (γ=-0.93, p=0.0002), while SMAD4 is known to interact with ANDR (AR), which might be involved in differential regulation of the AR transactivation (Kang et al. 2002). We predicted that TFE2 interact with PTF1A for predicting Tn (γ=5.22, p=3.69e-20), Tcm (γ=3.247, p=2.84e-29), Tem(γ=2.40, p=1.86e-13), and Temra (γ=-5.54, p=1.89e-68), while it has been reported that SMAD4 physically interacted with PTF1A and play a crucial role in blocking signal pathways of BMP-4, TCF-β 1 and activin A (Shimamoto et al. 1997). We predicted that HMGA2 interact with SMAD4 for predicting Tn (γ=-0.41, p=0.026), Tcm (γ=-2.24, p=2.84e-13) and Temra (γ=0.90, p=8.77e-05), while it is known that HMGA2 interacts with SMAD3/SMAD4 to regulate SNAIL1 gene expression (Thuault et al. 2008). The predicted interactions between known and unknown motifs as well as between unknown motifs are likely to be novel interactions, in particular those with strong and highly significant interactions, such as the interactions between M-Motif-71 and H3K4me1-30 that shows a negative interaction for predicting Tn (γ=-4.17, p=2.75e-274), Tcm(γ=-2.88, p=3.40e-115) and Tem (γ=-2.28, p=2.36e-100), but a positive interaction for predicting Temra(γ=3.78, p=2.09e-63). Similar patterns of interactions are seen in the models of the other five histone markers (Supplementary Figures 4 - 8).

**Figure 10.**
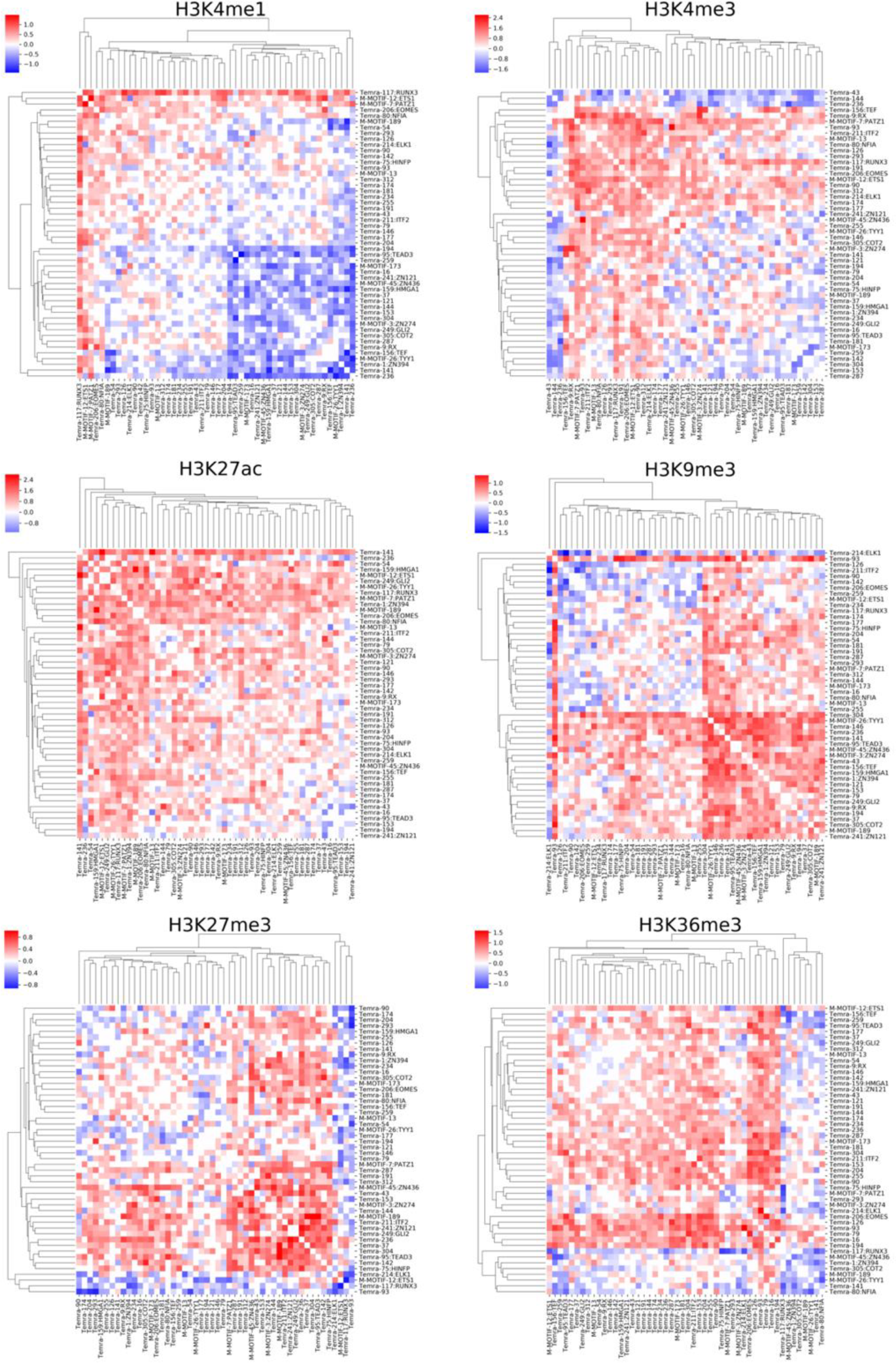
Interactions between each pair of the top 50 learned motifs on the predictions of the histone markers in the Temra cell model.

**Figure 11.**
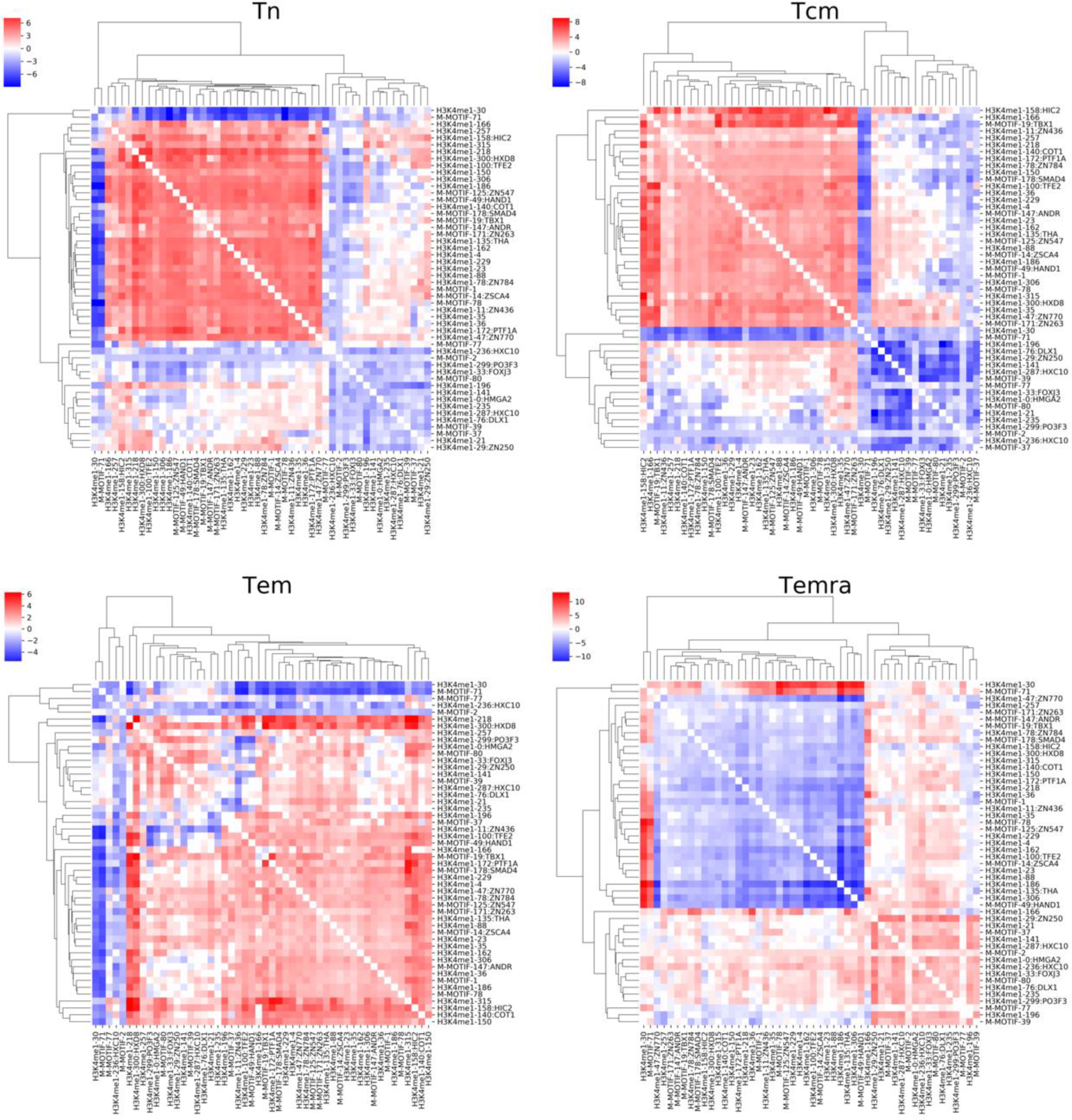
Interactions between each pair of the top 50 learned motifs on the predictions of the cell types in the H3K4me1 model.

## DISCUSSION

CNNs have been proved a powerful strategy to predict epigenomic features including TF binding (Alipanahi et al. 2015), DNase accessibility (Kelley et al. 2016), DNA methylation (Angermueller et al. 2017) and histone modifications (Zhou and Troyanskaya 2015). One of the advantages of CNNs, which other machine-learning method often lack, is their ability to automatically learn the features of the objects through the filters in the convolutional layers. However, how to interpret the learned features in the context of biological systems is still a challenging task (Wainberg et al. 2018). In the case of epigenomic analysis, these features include sequence determinants that define the unique patterns of epigenetic modifications in different cell types. Although efforts have been made to interpret the filters in CNN models in biological contexts (Kelley et al. 2016; Shrikumar et al. 2017), a single mixed model used in these studies to predict multiple labels each for a specific epigenetic marker in a specific cell type lacks the power of comparative analyses. To overcome this limitation and facilitate interpreting CNN models, we developed two types of CNN models to capture different features of sequences with various histone modifications in different cell types: 1) the cell type model for predicting patterns of various histone modifications in a cell type, and 2) the histone marker model for predicting various cell types based on a histone marker. In this way, by comparing the motifs earned in different cell type models, we could predict the common and unique motifs that specify unique patterns of various histone modifications in a cell type; and by comparing the motifs learned in different histone marker models, we could predict the common and unique motifs that determine different patterns of the same histone marker in different cell types. Furthermore, the models enable us to evaluate the impacts of learned motifs and heir interactions on the prediction accuracy, thereby predicting roles of each motifs in specifying histone modification and determining cell types.

To validate this strategy, after demonstrating the superior performance of both the cell models and histone marker models on an ENCODE dataset, we applied them to a dataset derived from four well-studied CD4+ T cell types in humans, i.e., Tn, Tcm, Tem and Temra, with high accuracy. A large portion of our predicted motifs resemble those of TFs that are known to play crucial roles in T cell development, while the remaining ones could be novel motifs of unknown TFs participating in T cell differentiation. By comparing the motifs learned in different cell models, we predicted that the unique patterns of various histone modifications in each cell type are largely determined by quite a unique set of motifs (Figure 5B and C), and at the same time, the number of common motifs shared by two cell models reflect linear lineage relationships of the four CD4+ T cell types (Figure 5G), which is consistent with the results based on DNA methylation, DNase hypersensitivity and transcription patterns in the earlier study that produced the datasets used in our analysis. Furthermore, by comparing the motifs learned in different histone marker models, we predicted that different patterns of the same histone markers in different cell types are largely determined by quite a unique set of motifs (Figure 5B and C), while the number of common motifs shared by two histone models reflect their co-modification and exclusiveness natures (Figure 5H). All these results suggest that at least of our predicted motifs are likely to be authentic, and play roles in T cell differentiation. Moreover, by computing the impact scores of the learned motifs, we further predicted the specific roles of each learned motif in determining the patterns of various histone marker modifications in a cell (Figure 6 A and C), or different patterns of the same histone modification in different cells (Figure 6B and D). Finally, by computing an interaction score, we predicted the interactions of the cognate TFs of the learned motifs in either the cell models or histone marker models. Many of these predictions have experimental supports. Thus, our results support the hypothesis that sequences ultimately determine the unique epigenomes of different cell types during the course of cell differentiation in a stepwise manner. Therefore, the motifs learned in our CNN models are highly interpretable and may provide insights into the underlying molecular mechanism of establishing the unique histone marker modifications in different cell types.

## MATERIALS AND METHODS

### The datasets

**Human embryonic stem cells datasets** We downloaded from the Roadmap Epigenomics Project (Roadmap Epigenomics et al. 2015) the ChIP-seq datasets for six histone markers H3K4me1, H3K4me3, H3K27me3, H3K27ac, H3K9me3 and H3K36me3 in human embryonic stem cells (H1) and in four cell types derived from H1, including trophoblast-like (TBL), mesendoderm (ME), mesenchymal (MSC) and neural progenitor (NPC) cells.

**Human CD4+ T cells datasets** We downloaded from European Genome-Phenome Archive the ChIP-seq datasets for six histone markers H3K4me1, H3K4me3, H3K27me3, H3K27ac, H3K9me3 and H3K36me3 in four different human CD4+ T cell types native T (Tn), central memory T (Tcm), T effector memory (Tem), and CD4+ terminally differentiated CD45RA+ memory T (Temra) cells (Durek et al. 2016).

### Peak calling and filtering

Peak calling To identify genome regions that are dominant with different histone modification markers, tight and broad histone modification peaks were called (Whitaker et al. 2015) (8). The tight peaks including H3K27ac, H3K4me1 and H3K4me3 are typically < 1 kbp. The broad peaks including H3K27me3, H3K36me3 and H3K9me3 are typically > 1 kbp. The tight peaks were called using MACS2 (Zhang et al. 2008) as follows: macs2 callpeak -t bam/tagAlign file -n name -c control file –outdir output dir -g hs -q 0.05 –nomodel – extsize fragment length

The broad peaks were called using MACS2 as follows: macs2 callpeak -t bam/tagAlign file -n name -c control file –outdir output dir -g hs –broad –broad-cutoff 0.1 –nomodel –extsize fragment size

The fragment sizes were estimated using phantompeakqualtools.

### Peak filtering and merging

We discarded peaks whose −*log*10(*qvalue*) is less than 2 or whose length is greater than 10,000 bp for their low quality or too long length. We also removed the peaks that overlap the blacklisted regions of the human genome (Consortium 2012), which are regions might show artificially high signal in all NGS experiments. To ensure only regions of high confidence are considered, we only used the intersection of at least two replicates when possible. We extracted and merged the peaks using BedTools (Quinlan and Hall 2010), and used the CRCh37/hg19 genome assembly for all the analyses.

### Convolutional neural networks

Convolutional neural networks (CNNs) are a type of feed-forward artificial neural networks. A CNN consists of an input layer, multiple convolutional layers and fully connected layers and an output layer. A convolutional layer convoluted with input signal using a specified number of kernels or filters. A pooling layer and a normalization layer are usually after convolutional layers to do down-sampling and prevent internal covariate shift (Ioffe and Szegedy 2015), respectively. Sequential alternating convolutional layer and pooling layer are frequently used to learn spatial structure of objects hierarchically. The fully connected layer can integrate the lower-level spatial structural information from the lower-level layers into higher-level spatial information. The output layer uses a sigmoid function to represent the final prediction as probability.

Our CNN models are made of a stack of three units each consisting a convolutional layer, a pooling layer and a batch normalization layer, followed by a fully connected layer and an output layer. We applied a Rectified linear transform as the activation function after a convolution layer, which could help to prevent vanishing gradient problem (Nair et al.; Lecun et al. 1998):

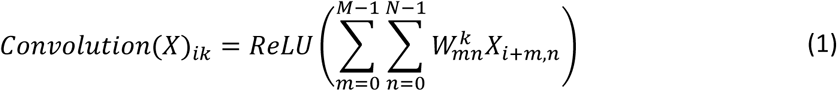

Where X is the input, i the index of the output position and k the index of the kernel.

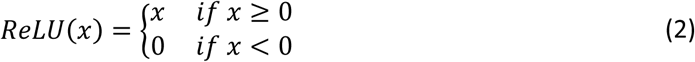

To decease internal covariate shift and accelerate training, we applied batch normalization layer after the convolutional layer (Ioffe and Szegedy 2015). Furthermore, we applied a max pooling layer after the batch normalization layer, which extract the maximum activation value from each of receptive field at the prior layer. Three convolutional layers contain 320, 300, 300 kernels, respectively, and the fully connected layer has 1,000 units with a sigmoid activation function feeding into the output layer. We implemented the CNN models using Theano (Team 2016) and Lasagne (Dieleman et al. 2015).

### Data representation

To prepare the input to the deep CNN models, we parsed the whole genome into 200-bp bins. For each bin, we label it with the corresponding histone marker (cell type model) or cell types (histone model) as 1 if the overlapping portion between the bin and a peak is above a threshold. To achieve the best prediction results, we tested different thresholds of 0.5, 0.6, 0.7, 0.8 and 0.9, and the threshold with the highest accuracy was used for the final analysis. We discarded those 200-bp bins that have no overlap with any histone modifications. We then extended the 200-bp bin into 1,000-bp sequence centered on the midpoint of the 200-bp bin. Each extended 1,000-bp sequence was represented by a 1,000 × 4 binary matrix, each column is one hot vector to represent the presence or absence of A, C, G, T at each nucleotide position, if a nucleotide position is N in the genome, we use [0.25, 0.25, 0.25, 0.25].

### Model training, validation and evaluation of prediction performance

We split dataset into a training dataset, a validation dataset and a test dataset with a ratio about 2:1:1, the objective function is binary cross entropy. Then we applied a stochastic gradient descent to minimize the objective function by updating all model parameters using RMSprop updates with learning rate 0.001 on minibatch (Hinton et al.). To avoid overfitting, we applied l1, l2 regularization terms and early stopping strategy. To keep the filters free to grow based on input sequences, we only applied l1, l2 regularization terms to the fully connected layer. To quickly choose the best set of hyperparameters of the models as Table 1, we used parallel random search. We applied l1, l2, and maximum epochs as shown in Table 1:

**Table 1.**
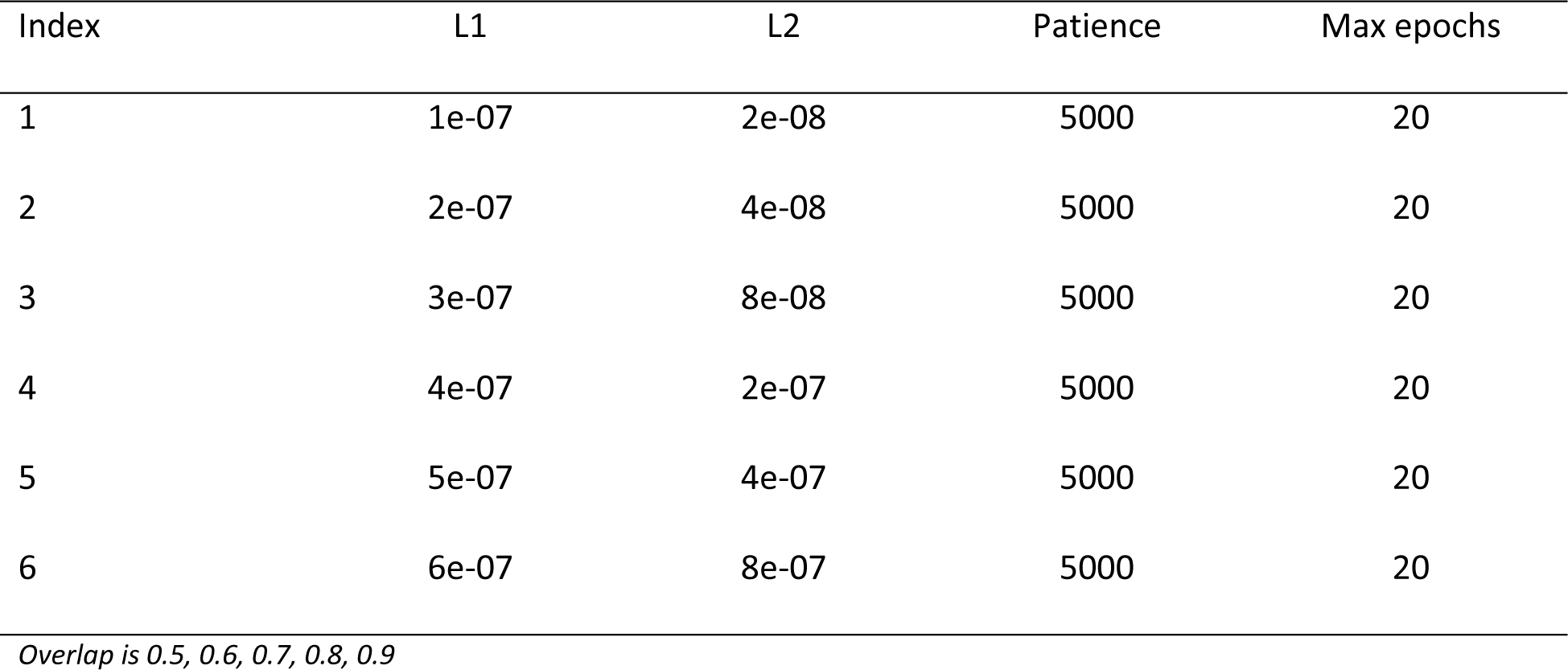
Hyper-parameter Configuration

We performed the receiver operating characteristic (ROC) curve analysis and used the area under the curve (AUC) to evaluate the performance of the models. We also define the accuracy of a model as follows:

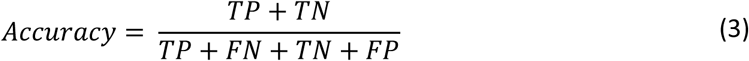

### Interpretation of the kernels/filters in the first convolutional layer

The first convolutional layer of models scans the DNA sequences with its kernel or filters to capture the composition of k-mers motifs that that differentiate modified and unmodified DNA sequences, thus potentially correspond to the binding motifs of TF or chromatin remodeling proteins that lead to the specific modifications at the loci. To reveal such possibility, we constructed a position weight matrices (PWMs) for each filter by extracting k-mers in the test dataset, which had a score against the filter greater than a threshold defined as,

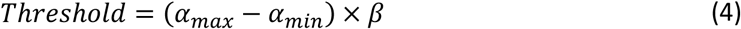

Where α_*max*_ and α_*min*_ are the maximum and minimum activations for a k-mer across all sequences in the test dataset, respectively, and β is a ration constant. For each filter, we evaluated β ranging from 0.3 to 0.8, and chose the resulting PWM with the highest information content. We discarded resulting PWMs with 0 information content. To evaluate the impact of a filter on the model’s prediction, we nullified forward information of the filter by setting its output as the its mean output over all nucleotides of all sequences in the test dataset (Kelley et al. 2016), and quantified each filter’s impact as sum of square of the difference of the prediction probability in the test dataset before and after the nullification as follows,

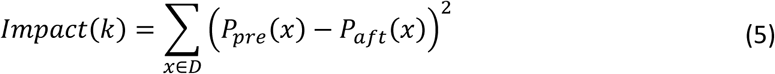

Where D is the test dataset, *P_pre_*(*X*) is the original prediction probability, *P_aft_*(*X*) the prediction probability after nullifying the filter k.

### Motif conservation analysis

We used Fimo (Grant et al. 2011) to scan sequences for binding sites of each motif as follows: fimo –parse-genomic-coord –thresh 1e-5 –bgfile fasta file background model –oc output_folder motifs_meme target_sequences

We used a 5th-order Markov model (Bailey et al. 2009) to generate the background sequences as follows: fasta-get-markov -m 5 -dna sequences background_model

We extracted the phastCons (Siepel et al. 2005) score for each position in each binding site, and calculated a conservation score for each motif as the mean PhastCons scores of all the binding sites of each motif learned in the models. To study the relationship between the impact of learned motifs and their conservation levels, we compute the Pearson correlation coefficient between them, and tested the null hypothesis of non-correlation using two-tailed p-values,

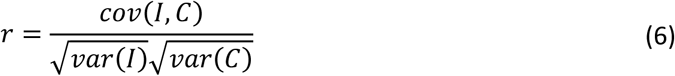

Where I, C are the impact and phastCons scores of motifs, respectively, and r Pearson correlation coefficient.

### Merging highly similar motifs

To merge similar motifs learned in all the cell and histone marker models, we compared each motifs with all other motifs using TOMTOM (Gupta et al. 2007), and constructed a graph by connecting two motifs if they are a pair of bidirectional best hits with a minimum overlap of 7 bps and E value < 0.1. We then cut the network into components using Networkx (Hagberg et al. 2008). We consider each resulting component as a unique motif. Some components are singletons containing a single original motif, while others are formed by merging multiple highly similar original motifs. To find the PWM for the merged motifs, we performed motifs finding on the merged binding sites using ProSampler (Li et al. 2018).

### Prediction of interactions between cognate TFs of learned motifs

To predict possible interactions between the cognate TF of the learned motifs, we applied a linear model to the changes in the prediction probability for random selected 2,000 sequences after the two motifs were simultaneously nullified, defined as:

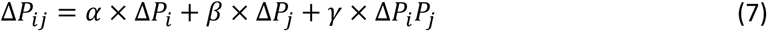

Where Δ*P_ij_* is sum square of changes in prediction probability after simultaneously nullifying motifs i and j, Δ*P_i_* and Δ*P_j_* are sum square of changes in prediction probability after nullifying motifs i and j separately.

## ACKNOWNLEDGES

The authors would like to thank Dr. Meng Niu for assisting in preprocessing the original ChIP-seq datasets, and members in the Su Lab for discussion. We also thank Dr.Polansky for providing the T cell Epigenome datasets. The work was partially supported by US National Science Foundation (DBI- 1661332) and NIH (R01GM106013) to ZS.

